# Gymnosperm FLS2 orthologues mediate conserved flg22 perception and divergent responses to flg22 variants

**DOI:** 10.64898/2026.04.10.717846

**Authors:** Hongju Xiao, Xiaodie Huo, Ling Wu, Lichi Zhong, Qiang Cheng

## Abstract

- Whether gymnosperm FLAGELLIN-SENSING 2 (FLS2) orthologues are functional receptors and whether their flg22-recognition spectra have already diversified remain unclear, despite the central role of FLS2 in flagellin perception in angiosperms.
- Here, we identified two gymnosperm FLS2 orthologues, GbFLS2 from *Ginkgo biloba* and PtFLS2 from *Pinus tabuliformis*, and analysed their function using transient and stable expression in the *Nicotiana benthamiana* fls2 mutant, in planta cross-linking assays, AlphaFold3 modelling and structure-guided mutagenesis.
- Both receptors restored flg22^Pst^-triggered reactive oxygen species production and MAPK activation, physically associated with flg22^Pst^, and required conserved residues for flg22^Pst^ recognition. In stable transgenic plants, both receptors mediated flg22^Pst^-triggered PTI outputs and flg22^Pst^-induced resistance to *Pseudomonas syringae*. PtFLS2 additionally mediated responsiveness to flg22^Rso^ and enhanced resistance to *Ralstonia solanacearum*, whereas GbFLS2 retained a typical flg22^Pst^-recognition profile.
- These findings provide direct evidence that gymnosperm FLS2 orthologues can function as bona fide flagellin receptors in a heterologous angiosperm background, and further suggest that diversification in flg22 recognition had already emerged within gymnosperms.

## Introduction

Plants rely on cell-surface pattern-recognition receptors (PRRs) to detect conserved microbial molecules and activate pattern-triggered immunity (PTI). Among the best-characterized PRRs, FLAGELLIN-SENSING 2 (FLS2) is a leucine-rich repeat receptor-like kinase (LRR-RLK) that recognizes the flagellin-derived epitope flg22. Upon ligand perception, FLS2 recruits the co-receptor BAK1 to form an activated receptor complex, in which flg22 acts as a molecular bridge stabilizing the FLS2 – BAK1 ternary complex and initiating downstream immune signalling (Heese et al., 2007; Chinchilla et al., 2007; Sun et al., 2013). Because its ligand, signalling outputs and activation mechanism are well defined, the FLS2 pathway has become a model system for studying receptor activation in plant immunity.

FLS2 homologues have been functionally characterized in diverse angiosperms, indicating that flg22 perception is broadly conserved in this lineage. At the same time, naturally occurring and engineered FLS2 variants can differ substantially in ligand-recognition range, including expanded responsiveness to polymorphic flg22 peptides (Li et al., 2025; Zhang et al., 2025). In parallel, the generation of a *Nicotiana benthamiana fls2* mutant has provided a useful heterologous platform for testing the flagellin-recognition spectrum of diverse FLS2 receptors (Wu et al., 2022). These studies raise two questions: whether FLS2-mediated flagellin perception is conserved beyond angiosperms, and whether diversification in flg22 recognition had already emerged in other seed plant lineages. However, despite the central role of FLS2 in angiosperm immunity, direct experimental evidence for functional FLS2 receptors outside angiosperms is still lacking. It therefore remains unknown whether non-angiosperm seed plants simply retain canonical flagellin perception or have already diversified in ligand recognition.

Gymnosperms are well suited for addressing these questions. Together with angiosperms, they comprise the two extant seed plant lineages, and phylogenomic analyses support their deep divergence on an evolutionary timescale of roughly 300 million years (Ran et al., 2018). Extant gymnosperms include ginkgo, cycads, gnetophytes and conifers. Their long independent evolutionary history makes them useful for testing whether an ancient immune receptor module has been conserved, diversified, or both across seed plants. In this context, *G. biloba*, the sole extant representative of Ginkgoales, and *P. tabuliformis*, a representative conifer species, provide a useful comparison across deeply separated gymnosperm branches. The availability of high-quality genome resources for both species now enables systematic identification and functional analysis of candidate immune receptors in these non-angiosperm lineages (Liu et al., 2021; Niu et al., 2022).

Here, using *Arabidopsis thaliana* FLS2 as a reference, we identified two gymnosperm FLS2 orthologues, GbFLS2 from *G. biloba* and PtFLS2 from *P. tabuliformis*. We asked whether these receptors are functional FLS2 orthologues, whether they retain conserved ligand-recognition features and dependence on the BAK1 co-receptor module, whether they remain functional in an angiosperm signalling background, and whether diversification in flg22 recognition had already emerged in gymnosperms. By combining transient complementation, stable transgenic analysis, in planta ligand-association assays, structural modelling, mutational validation and pathogen infection experiments, we show that these two gymnosperm FLS2 orthologues can function as bona fide flagellin receptors in a heterologous angiosperm background while exhibiting diversified flg22 recognition. These findings show that functional FLS2 receptors also occur in gymnosperms and that gymnosperms remain underexplored for FLS2 diversity.

## Materials and Methods

### Plant materials and growth conditions

*N. benthamiana fls2* mutant plants (Wu et al., 2022) were grown in a controlled growth chamber at 24°C under a 16 h light/8 h dark photoperiod with approximately 60% relative humidity. Four-week-old plants were used for transient expression, reactive oxygen species (ROS) burst assays, MAPK activation assays and in planta cross-linking assays. For seedling growth inhibition assays, seeds of homozygous T2 lines of *N. benthamiana fls2* expressing AtFLS2, GbFLS2 or PtFLS2, together with the *fls2* mutant control, were surface sterilized, germinated on solid half-strength Murashige and Skoog (1/2 MS) medium for 1 week, and then transferred to liquid 1/2 MS medium containing the indicated flg22 peptides for an additional week. Young leaves of *G. biloba* and *P. tabuliformis* used for gene cloning were collected from plants growing on the campus of Nanjing Forestry University, Nanjing, China.

### Identification of gymnosperm FLS2 orthologues and plasmid construction

The amino acid sequence of *A. thaliana* FLS2 (AtFLS2) was used as a query to search the genome resources of *G. biloba* and *P. tabuliformis* by tBLASTn. Candidate loci were further evaluated based on predicted gene structure, domain composition and reciprocal BLAST analysis against the Arabidopsis proteome. The best-supported candidates were designated GbFLS2 and PtFLS2, respectively.

Using gene-specific primers (Table S1), full-length genomic fragments of *AtFLS2, AtEFR, GbFLS2* and *PtFLS2*, encompassing the complete coding sequences including introns, were amplified from plant genomic DNA and cloned using the ClonExpress II One Step Cloning Kit. For transient expression and in planta cross-linking assays, the genomic fragments were inserted into pH35GG-Km, in which expression is driven by the CaMV 35S promoter and the receptor proteins carry an in-frame C-terminal GFP tag. For stable transformation, corresponding untagged genomic constructs were generated in pH35GS-Km, because C-terminal tagging has been reported to compromise FLS2 function (Hurst et al., 2018).

### AlphaFold3 modelling and interface analysis

The extracellular domains of GbFLS2 and PtFLS2 were modelled in complex with flg22^Pst^ or flg22^Rso^ using the AlphaFold3 server. For each receptor–ligand combination, multiple independent predictions were generated using different random seeds. For most receptor–ligand combinations, 10 independent models were included in the final analysis. For the PtFLS2 – flg22^Rso^ complex, the analysis was extended to 16 independent models to further assess interface reproducibility. Hydrogen-bonding interactions and structural visualization were analysed in PyMOL. To quantify local interface features, a custom script was used to identify FLS2 residues within 5 Å of each residue of the 22-amino-acid flg22 peptide and to extract the corresponding inter-residue distances and predicted aligned error (PAE) values from the AlphaFold3 output files.

### Protein-blot analysis of in planta cross-linking assays

C-terminally biotinylated peptides, including flg22^Pst^-EDA-biotin (TRLSSGLKINSAKDDAAGLQIA), flg22^Rso^-EDA-biotin (QRLSTGLRVNSAQDDSAAYAAS) and flg22^Xcc^-EDA-biotin (QQLSSGKRITSASVDAAGLAIS), together with the corresponding unlabelled peptides, were synthesized by GenScript at >85% purity. In planta cross-linking assays were performed in *N. benthamiana fls2* leaves transiently expressing C-terminal GFP-tagged AtFLS2, AtEFR, GbFLS2 or PtFLS2 at 48 h after Agrobacterium infiltration. Biotinylated flg22 peptides (50 nM) were infiltrated in HEPES buffer (pH 7.5) containing 2 mM ethylene glycol bis (succinimidyl succinate) (EGS). For competition treatments, 100 μM of the corresponding unlabelled peptide was added together with the biotinylated peptide. Control leaves were infiltrated with the same buffer containing EGS but without peptide. After infiltration, leaves were incubated at room temperature for 20 min to allow in planta cross-linking. Leaf discs were collected and total proteins were extracted using Plant RIPA lysis buffer (Beyotime). GFP-tagged proteins were purified using Anti-GFP Affinity Gel (MCE). Immunoprecipitated proteins were separated by 12% SDS–PAGE and transferred onto Hybond ECL membranes. Receptor-associated biotinylated peptides were detected using alkaline phosphatase-conjugated streptavidin (Sangon Biotech), whereas immunoprecipitated GFP-tagged proteins were detected with an anti-GFP antibody (GenScript) followed by an alkaline phosphatase-conjugated secondary antibody.

Both signals were visualized with BCIP/NBT substrate solution (Chinchilla et al., 2006; Albert et al., 2015).

### Pathogen infection assays

For bacterial infection assays, *N. benthamiana fls2* mutant plants and transgenic lines expressing AtFLS2, GbFLS2 or PtFLS2 were used. For *Pseudomonas syringae* pv. *tomato* DC3000 Δ hopQ1-1 infection assays, plants were pre-treated with 1 μ M flg22^Pst^ for 12 h and then spray-inoculated with bacterial suspensions at 10^7^ CFU mL−1 in 10 mM MgCl_2_ containing Silwet L-77. Disease symptoms were recorded at 7 d post inoculation, and bacterial growth was quantified at 2 and 4 d post inoculation by plating serial dilutions of leaf extracts on King’s B medium containing rifampicin. For *Ralstonia solanacearum* strain TP2 infection assays, plants were inoculated by soil drenching. Bacterial suspensions were adjusted to OD600 = 0.5, and each pot was drenched with 50 mL inoculum. Wilt symptoms were scored using a 0 – 4 disease index, and plant survival was recorded at the indicated time points.

### Statistical analysis

Statistical analyses were performed using GraphPad Prism 10. Data are presented as mean ± SD unless otherwise indicated. Details of statistical tests and sample sizes are provided in the figure legends.

## Results

### GbFLS2 and PtFLS2 are functional gymnosperm FLS2 receptors that mediate canonical flg22^Pst^-triggered signalling in transient assays

Using *A. thaliana* FLS2 (AtFLS2) as a query, tBLASTn searches against the genome assemblies of *G. biloba* and *P. tabuliformis* identified one candidate FLS2-like locus from each species. Full-length genomic fragments encompassing the entire coding regions, including introns, were subsequently obtained by homology-based cloning and were designated *GbFLS2* (GenBank accession no. PZ262359) and *PtFLS2* (GenBank accession no. PZ262360), respectively. GbFLS2 and PtFLS2 encode typical plasma membrane-localized receptor-like kinases containing an N-terminal signal peptide, an extracellular leucine-rich repeat (LRR) domain, a single transmembrane domain and an intracellular kinase domain. Both proteins contain 28 extracellular LRRs, as reported for most previously characterized FLS2 receptors, and harbour a non-RD kinase domain, a hallmark of many immune receptor kinases (Fig. S1). In a maximum-likelihood phylogenetic analysis of representative FLS2 proteins, GbFLS2 and PtFLS2 grouped within the FLS2 clade and formed a well-supported sister branch to angiosperm FLS2 receptors, clearly separated from other representative members of the RLK XII family included for comparison (Fig. 1A). An expanded phylogenetic analysis including 112 angiosperm FLS2 sequences further supported the placement of GbFLS2 and PtFLS2 within the FLS2 clade and their separation from angiosperm FLS2 receptors (Fig. S2).

**Figure 1.**
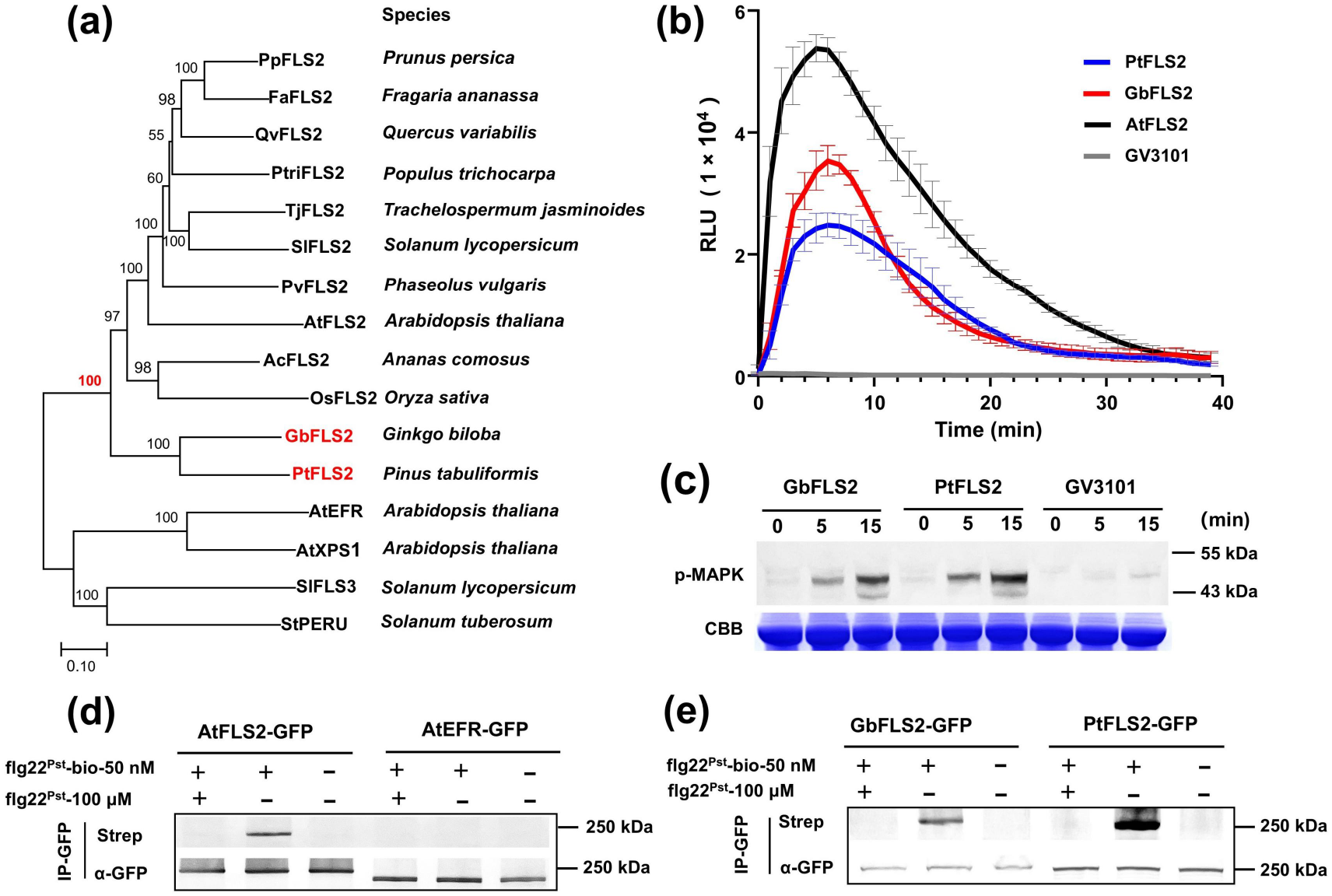
GbFLS2 and PtFLS2 are functional gymnosperm FLS2 receptors that mediate flg22^Pst^-triggered signalling and associate with flg22^Pst^ in planta. (A) Maximum-likelihood phylogenetic tree of GbFLS2 and PtFLS2 together with representative functionally validated angiosperm FLS2 receptors. Four known PRRs from the RLK XII family were included for comparison. Bootstrap support values were calculated from 1000 replicates and are shown at the corresponding nodes. Scale bar, substitutions per site. (B) Reactive oxygen species (ROS) burst triggered by flg22^Pst^ in *N. benthamiana fls2* leaf discs transiently expressing AtFLS2, GbFLS2 or PtFLS2. Leaf discs were treated with 200 nM flg22^Pst^. GV3101 carrying empty vector was used as a negative control. ROS was measured by a luminol-based assay. Data are presented as mean ± SD (n = 6 leaf discs). Data are shown from one representative experiment repeated three times with similar results. (C) flg22^Pst^-induced MAPK activation in *N. benthamiana fls2* leaf discs transiently expressing GbFLS2 or PtFLS2. Leaf discs were treated with 1 μM flg22Pst for the indicated times, and phosphorylated MAPKs were detected by immunoblot using an anti-phospho-p44/42 MAPK antibody. Coomassie brilliant blue (CBB) staining is shown as a loading control. (D) Protein-blot analysis of in planta cross-linking assays validating flg22^Pst^ association with AtFLS2-GFP. *N. benthamiana fls2* leaves expressing AtFLS2-GFP or AtEFR-GFP were treated with C-terminally biotinylated flg22^Pst^ (flg22Pst-bio, 50 nM) in the presence or absence of excess unlabelled flg22^Pst^ (100 μM), followed by in planta cross-linking with EGS. GFP-tagged receptors were immunoprecipitated with GFP-Trap beads, and receptor-associated flg22^Pst^-bio was detected with alkaline phosphatase-conjugated streptavidin followed by BCIP/NBT staining (Strep). Immunoprecipitated GFP-tagged proteins were detected with an anti-GFP antibody. (E) Protein-blot analysis of in planta cross-linking assays showing association of flg22^Pst^ with GbFLS2-GFP and PtFLS2-GFP in *N. benthamiana fls2*. Samples were analysed as in (D).

We next asked whether these two gymnosperm homologues could mediate flg22-triggered immune signalling. Transient expression of GbFLS2 or PtFLS2 in the *N. benthamiana fls2* mutant restored responsiveness to flg22^Pst^, as shown by a rapid oxidative burst after peptide treatment (Fig. 1B). The ROS kinetics of GbFLS2 and PtFLS2 were similar to those of AtFLS2. All three receptors produced a rapid and transient burst that peaked at approximately 5–7 min after treatment, although AtFLS2 consistently gave a stronger peak. Consistent with the ROS assay, flg22^Pst^ treatment also triggered rapid MAPK activation in leaf discs expressing GbFLS2 or PtFLS2 (Fig. 1C). Using an anti-phospho-p44/42 MAPK antibody, phosphorylated MAPK signals in the 43–55 kDa range were detectable as early as 5 min after treatment and increased by 15 min.

To determine whether GbFLS2 and PtFLS2 physically associate with flg22^Pst^ in planta, we performed protein-blot analysis of in planta cross-linking assays using C-terminally biotinylated flg22^Pst^. Biotinylated flg22^Pst^ was infiltrated into *N. benthamiana fls2* leaves expressing GFP-tagged receptors, followed by in planta cross-linking with EGS and immunoprecipitation with GFP-Trap beads. Receptor-associated flg22^Pst^-bio was then detected with alkaline phosphatase-conjugated streptavidin followed by BCIP/NBT staining. We first validated this assay using AtFLS2-GFP as a positive control and AtEFR-GFP as a negative control. Biotinylated flg22^Pst^ was readily detected in AtFLS2-GFP immunoprecipitates, but not in AtEFR-GFP immunoprecipitates, and the AtFLS2-associated signal was abolished by competition with excess unlabelled flg22^Pst^ (100 μ M), indicating that the assay can monitor specific FLS2 – flg22 association in planta (Fig. 1D). We then applied the same assay to GbFLS2-GFP and PtFLS2-GFP. In both cases, receptor-associated flg22^Pst^-bio was detected as a specific band of approximately 250 kDa after GFP immunoprecipitation, whereas the signal was strongly reduced or undetectable in the presence of excess unlabelled flg22^Pst^ (Fig. 1E). Together with their placement within the FLS2 clade and their ability to mediate flg22^Pst^-induced ROS burst and MAPK activation, these results support the conclusion that GbFLS2 and PtFLS2 are bona fide gymnosperm FLS2 receptors.

### GbFLS2 and PtFLS2 recognize flg22^Pst^ through conserved interaction interfaces

To define the structural features underlying flg22^Pst^ recognition by GbFLS2 and PtFLS2, we generated AlphaFold3 (AF3) models of the GbFLS2 – flg22Pst – NbSERK3B and PtFLS2 – flg22Pst – NbSERK3B ternary complexes, using NbSERK3B as a representative *N. benthamiana* BAK1 orthologue. Both complexes yielded high-confidence predictions across 10 independent runs, with consistently high ipTM values (GbFLS2, 0.883 ± 0.007; PtFLS2, 0.902 ± 0.005) (Fig. S3). We then examined the predicted FLS2 – flg22 interface, which was highly reproducible across replicate models. At the predicted GbFLS2–flg22^Pst^ interface, nine residues of flg22^Pst^ formed recurrent hydrogen-bonding interactions with 13 GbFLS2 residues in 7–10 of the 10 predictions, whereas at the predicted PtFLS2–flg22^Pst^ interface, seven residues of flg22^Pst^ formed recurrent hydrogen-bonding interactions with 10 PtFLS2 residues in 9–10 of the 10 predictions. These recurrent contacts were also associated with low mean pairwise PAE values (2.5–5.52 Å), indicating that the predicted local interface was reliable. Most of these interface residues mapped to the LRR6–LRR16 region in both receptors and were represented by positionally conserved residues with similar physicochemical properties (Fig. 2A,B; Fig. S1; Tables S2,S3).

**Figure 2.**
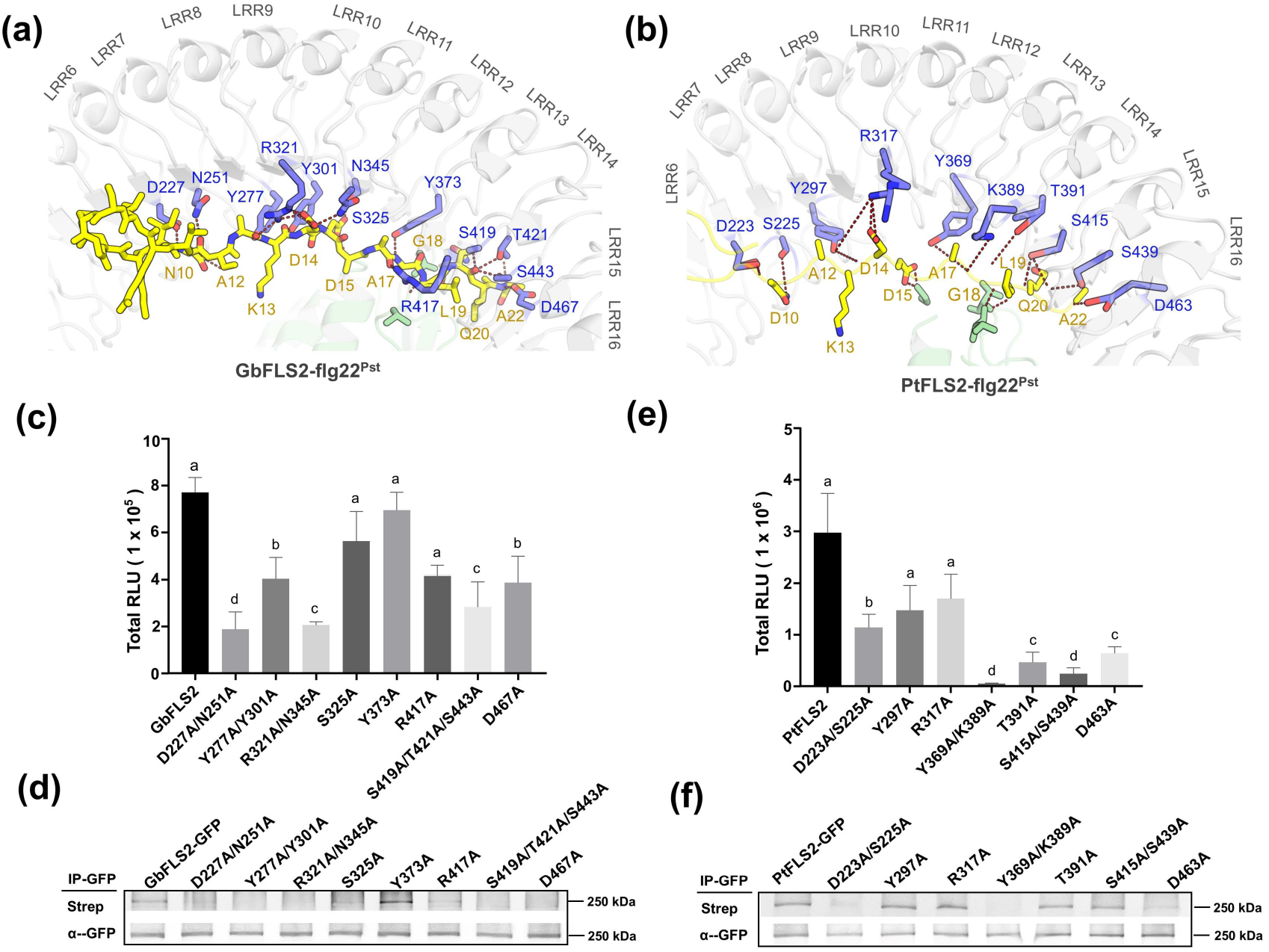
Structure-guided mutational analysis defines the flg22^Pst^-binding interfaces of GbFLS2 and PtFLS2. (A, B) Local views of the predicted interaction pockets in the GbFLS2–flg22^Pst^ and PtFLS2– flg22^Pst^ complexes, respectively, based on AF3 models. flg22^Pst^ residues are shown in yellow and receptor residues in blue. Predicted hydrogen bonds are indicated by dashed lines, and interacting residue pairs are labelled. (C, E) flg22Pst-induced reactive oxygen species (ROS) production in *N. benthamiana fls2* leaf discs transiently expressing the indicated GbFLS2 or PtFLS2 mutants. Leaf discs were treated with 200 nM flg22^Pst^. ROS was measured by a luminol-based assay, and total ROS accumulation is shown as integrated RLU values over 30 min. Data are presented as mean ± SD (n = 6 leaf discs). Different letters indicate statistically significant differences as determined by one-way ANOVA followed by Tukey’s multiple-comparison test (P < 0.05). (D, F) Protein-blot analysis of in planta cross-linking assays showing association of flg22^Pst^ with the indicated GbFLS2 or PtFLS2 mutants in *N. benthamiana fls2*. Samples were analysed as in Fig. 1D.

To test the functional importance of these predicted interface residues, we generated a series of structure-guided substitutions in GbFLS2 and examined their effects on flg22^Pst^-induced ROS production and ligand association in the in planta cross-linking assay. In GbFLS2, alanine substitutions D227/N251, Y277/Y301, R321/N345, S419/T421/S443 and D467 all markedly reduced the ROS response, whereas substitutions at S325, Y373 and R417 had little or no significant effect on ROS output (Fig. 2C). In the in planta cross-linking assay, most GbFLS2 substitutions likewise reduced receptor-associated flg22^Pst^-bio, whereas Y373A retained detectable ligand association (Fig. 2D). Thus, in GbFLS2, the effects of predicted interface substitutions on ligand association and ROS output were broadly consistent.

We next applied the same strategy to PtFLS2. In PtFLS2, substitutions D223A/S225A, Y369A/K389A and D463A reduced both flg22^Pst^-induced ROS production and receptor-associated flg22^Pst^-bio, whereas T391A and S415A/S439A reduced ROS output but retained detectable ligand association in the in planta cross-linking assay (Fig. 2E, F). These results support the predicted PtFLS2 – flg22^Pst^ interface, although the effects of the substitutions on ligand association and signalling were less consistent than in GbFLS2.

### PtFLS2 displays diversified flg22 recognition and mediates perception of flg22^Rso^

To compare the flg22 recognition spectra of the two gymnosperm receptors, we transiently expressed GbFLS2 or PtFLS2 in the *N. benthamiana fls2* plant and examined their responses to flg22 peptides from *Pseudomonas* species and four well-known flg22 variants that typically fail to trigger strong responses through angiosperm FLS2 receptors. GbFLS2 showed a response profile consistent with that of a typical FLS2 receptor and did not respond to the four tested flg22 variants (Fig. 3A). By contrast, PtFLS2 responded not only to the Pseudomonas-derived flg22 peptides, but also to flg22^Rso^ from *Ralstonia solanacearum* (Fig. 3B). In addition, PtFLS2 reproducibly showed a weak inducible response to flg22^Xcc^ (Fig. 3B; Fig. S3).

**Figure 3.**
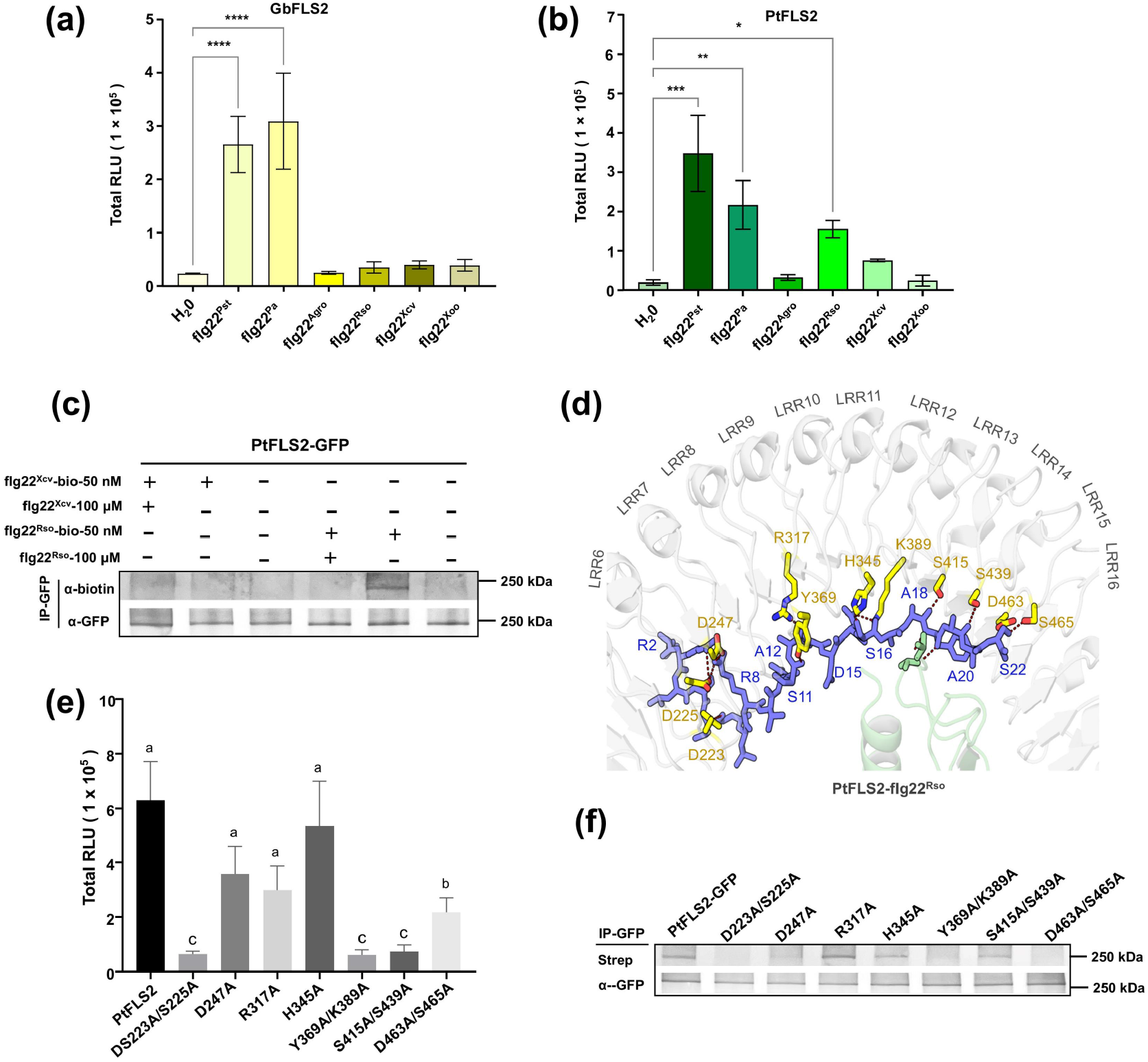
PtFLS2 displays diversified flg22 recognition and mediates perception of flg22^Rso^. (A, B) Reactive oxygen species (ROS) burst triggered by flg22 peptides in *N. benthamiana fls2* leaf discs transiently expressing GbFLS2 or PtFLS2. Leaf discs were treated with the indicated peptides at 200 nM. flg22^Pst^, *Pseudomonas syringae* pv. *tomato*; flg22^Pae^, *Pseudomonas aeruginosa*; flg22^Agro^, *Agrobacterium tumefaciens*; flg22^Rso^, *Ralstonia solanacearum*; flg22^Xcc^, *Xanthomonas campestris* pv. *campestris*; flg22^Xoo^, *Xanthomonas oryzae* pv. *oryzae*. (C) Protein-blot analysis of in planta cross-linking assays showing specific association of PtFLS2-GFP with flg22^Rso^, but not flg22^Xcc^, in *N. benthamiana fls2*. Samples were analysed as in Fig. 1D. (D) Local view of the predicted interaction pocket in the PtFLS2–flg22^Rso^ complex based on AF3 models. flg22^Rso^ residues are shown in yellow and receptor residues in blue. Predicted hydrogen bonds are indicated by dashed lines, and interacting residue pairs are labelled. (E) flg22^Rso^-induced reactive oxygen species (ROS) production in *N. benthamiana fls2* leaf discs transiently expressing the indicated PtFLS2 alanine-substitution mutants. (F) Protein-blot analysis of in planta cross-linking assays showing association of flg22^Rso^ with the indicated PtFLS2 mutants in *N. benthamiana fls2*. For (A), (B) and (E), ROS was measured by a luminol-based assay, and total ROS accumulation is shown as integrated RLU values over 30 min. Data are presented as mean ± SD (n = 6 leaf discs). In (A) and (B), asterisks indicate significant differences relative to the H_2_O control as determined by Student ‘ s t-test. In (E), different letters indicate statistically significant differences as determined by one-way ANOVA followed by Tukey ‘ s multiple-comparison test (P < 0.05).

To determine whether the PtFLS2-mediated response to flg22^Rso^ reflects specific ligand association in planta, we next performed the in planta cross-linking assay using biotinylated flg22^Rso^ and flg22^Xcc^. Receptor-associated flg22^Rso^-bio was readily detected in PtFLS2-GFP immunoprecipitates, and this signal was abolished by competition with an excess of unlabelled flg22^Rso^ (Fig. 3C). By contrast, flg22^Xcc^-bio did not produce a comparable specific signal under the same conditions. These results show that PtFLS2 specifically associates with flg22^Rso^ in planta.

To explore the structural features associated with PtFLS2-mediated flg22^Rso^ perception, we generated AF3 models of the PtFLS2–flg22^Rso^ complex. These models yielded a relatively high overall confidence score (ipTM = 0.8525 ± 0.0093), but were less convergent than the corresponding PtFLS2 – flg22^Pst^ models, even after expanding the analysis to 16 independent models. Predicted hydrogen-bonding contacts showed lower reproducibility, being recovered in 7–15 of the 16 models, and pairwise PAE values at the putative interface were substantially higher (11.27–17.66 Å) (Fig. 3D; Table S4). Nevertheless, several PtFLS2 residues implicated in flg22^Pst^ recognition remained recurrently positioned near flg22^Rso^ across replicate models, whereas two additional residues were identified as candidate contact residues found only in the PtFLS2–flg22^Rso^ models.

To test the functional relevance of these residues, we generated a set of alanine substitutions in PtFLS2 and examined their effects on flg22^Rso^-induced ROS production and ligand association in the in planta cross-linking assay. Alanine substitutions in hydrogen-bonding residues shared between the flg22^Pst^ and flg22^Rso^ models severely impaired the ROS response to flg22^Rso^ (Fig. 3E), and two of these also abolished receptor-associated flg22^Rso^-bio (Fig. 3F). By contrast, substitutions of the two candidate flg22^Rso^-specific residues, D247A and H345A, did not noticeably affect either flg22^Rso^-induced ROS production or ligand association (Fig. 3E, F). Thus, our data support flg22^Rso^ perception by PtFLS2 and validate a subset of shared interface residues, but do not yet resolve the full set of flg22^Rso^-specific determinants.

### GbFLS2 and PtFLS2 mediate conserved PTI outputs in stable transgenic *N. benthamiana*

To determine whether gymnosperm FLS2 receptors remain functional in stable transgenic plants, we generated *N. benthamiana fls2* lines expressing untagged *35S::AtFLS2, 35S::GbFLS2* or *35S::PtFLS2*. Two independent transgenic lines were selected for each construct and used for subsequent analyses. All transgenic lines restored responsiveness to flg22^Pst^, as shown by a robust ROS burst after peptide treatment, whereas the *fls2* mutant showed no detectable response (Fig. 4A). By contrast, only the PtFLS2 transgenic lines responded strongly to flg22^Rso^, while the AtFLS2- and GbFLS2-expressing lines remained indistinguishable from the *fls2* mutant background (Fig. 4B).

**Figure 4.**
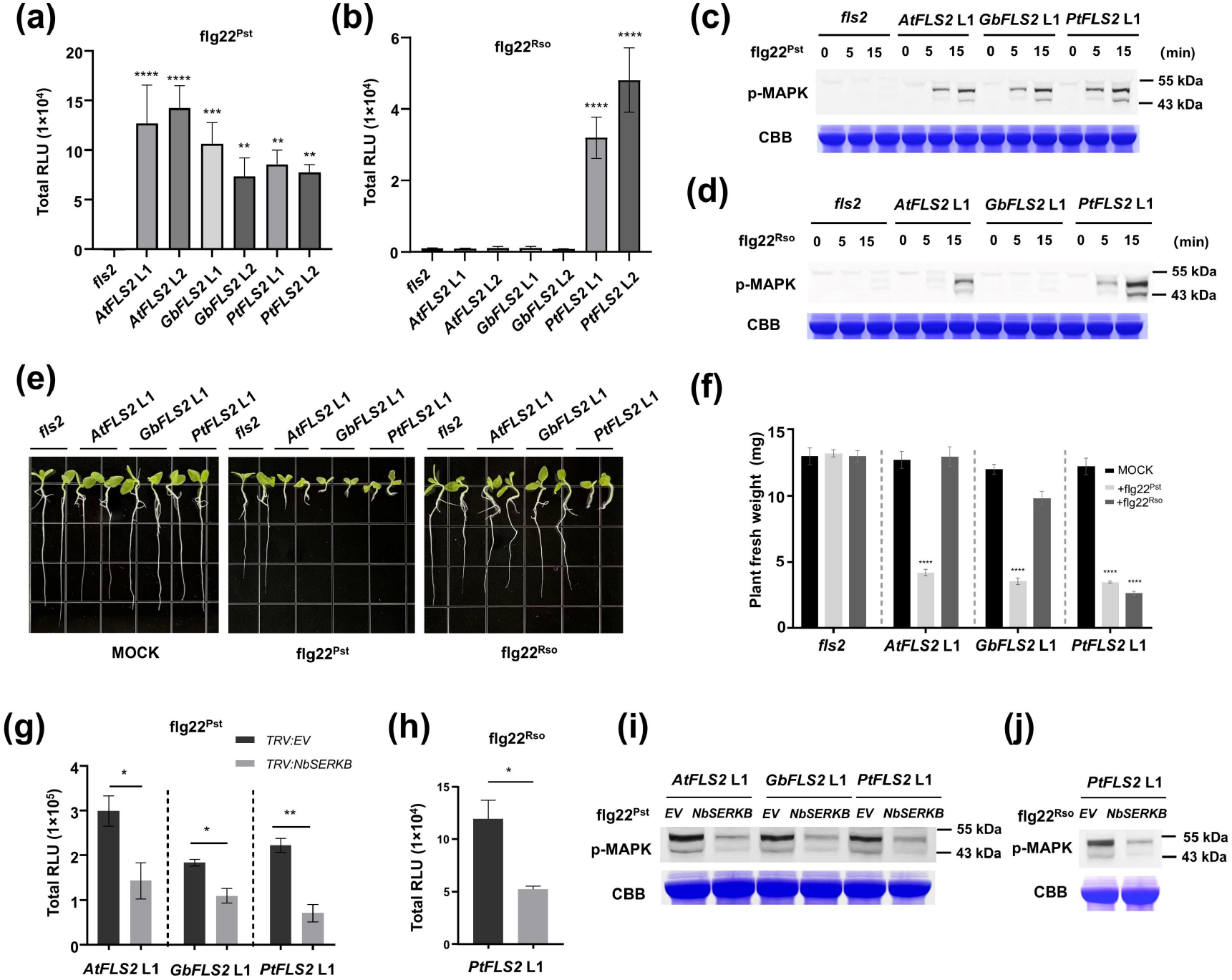
GbFLS2 and PtFLS2 mediate conserved PTI outputs in stable transgenic *N. benthamiana*. (A, B) Reactive oxygen species (ROS) burst triggered by flg22^Pst^ or flg22^Rso^ in *N. benthamiana fls2* and stable transgenic lines expressing AtFLS2, GbFLS2 or PtFLS2. Two independent transgenic lines were analysed for each construct. Leaf discs were treated with 200 nM flg22^Pst^ (A) or 200 nM flg22^Rso^ (B), and ROS production was measured by a luminol-based assay. Total ROS accumulation is shown as integrated RLU values over 30 min. Data are presented as mean ± SD (n = 6 leaf discs). Asterisks indicate significant differences relative to the *fls2* mutant as determined by Student’s t-test. (C, D) flg22-induced MAPK activation in representative L1 stable transgenic lines. Leaf discs were treated with 1 μ M flg22^Pst^ (C) or 1 μ M flg22^Rso^ (D) for the indicated times, and phosphorylated MAPKs were detected by immunoblot using an anti-phospho-p44/42 MAPK antibody. Coomassie brilliant blue (CBB) staining is shown as a loading control. Similar results for the corresponding L2 lines are shown in Fig. S4. (E) Seedling growth inhibition assay in representative L1 stable transgenic lines. Seedlings grown on solid medium for 1 week were transferred to liquid medium containing 5 μ M flg22^Pst^, 5 μ M flg22^Rso^ or mock medium without peptide, and cultivated for an additional week before imaging. Corresponding results for the L2 lines are shown in Fig. S4. (F) Fresh-weight measurements corresponding to the seedling growth inhibition assay shown in (E). Data are presented as mean ± SD (n = 12 seedlings). Asterisks indicate significant differences relative to the corresponding mock treatment as determined by Student’ s t-test. Corresponding results for the L2 lines are shown in Fig. S4. (G, H) Effects of *NbSERK3B* silencing on ROS production in representative L1 transgenic lines treated with 200 nM flg22^Pst^ (G) or 200 nM flg22^Rso^ (H). ROS was analysed as in (A, B). (I, J) Effects of *NbSERK3B* silencing on flg22-induced MAPK activation in representative L1 transgenic lines treated with 1 μ M flg22^Pst^ for 15 min (I) or 1 μ M flg22^Rso^ for 15 min (J). MAPK activation was analysed as in (C, D).

**Figure 5.**
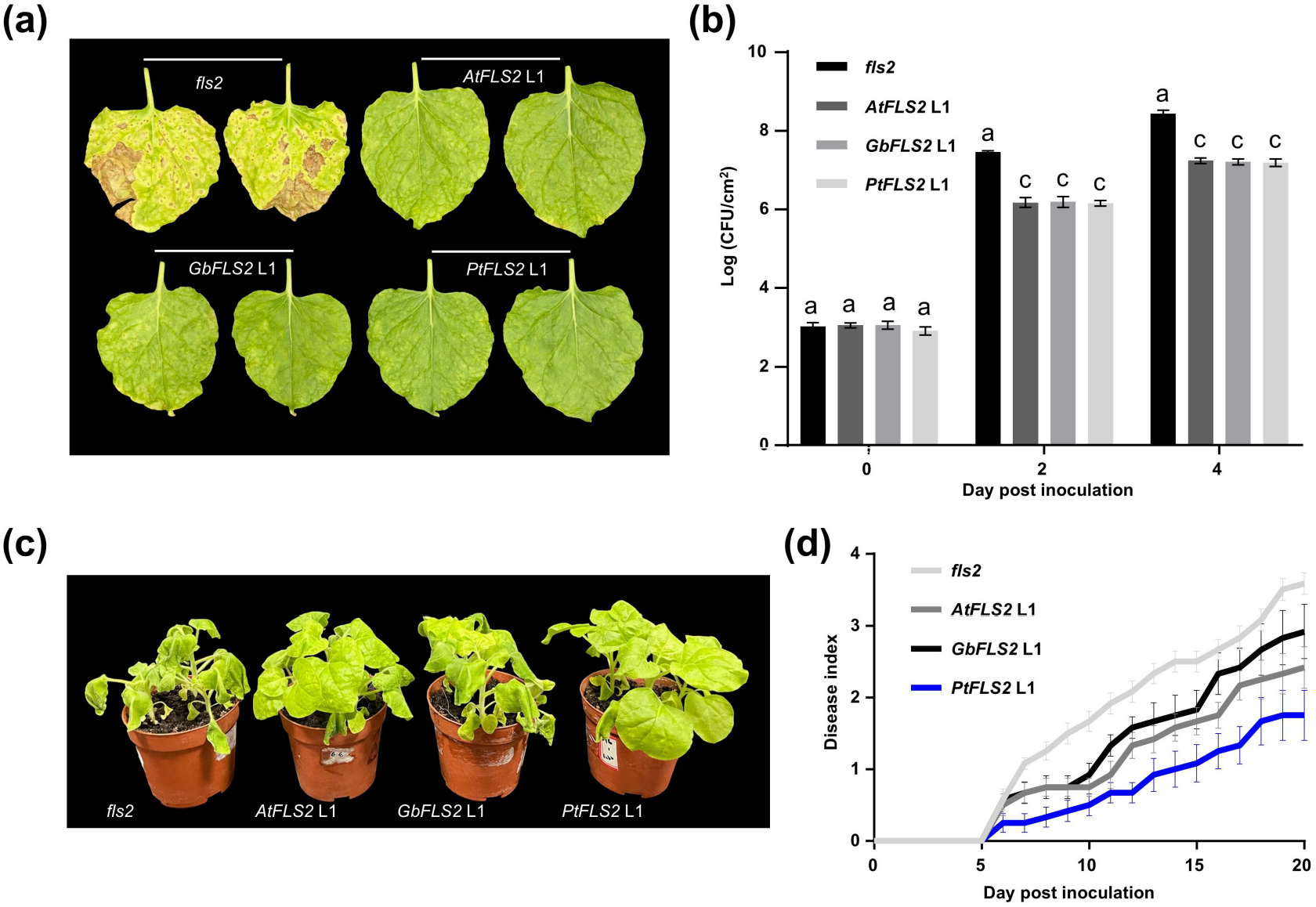
Stable expression of gymnosperm FLS2 receptors enhances antibacterial immunity in transgenic *N. benthamiana*. (A) Representative disease symptoms in *N. benthamiana fls2* and transgenic lines expressing AtFLS2, GbFLS2 or PtFLS2 after spray inoculation with *Pseudomonas syringae* pv. *tomato* DC3000 Δ hopQ1-1 following flg22^Pst^ pre-treatment. Plants were pre-treated with 1 μ M flg22^Pst^ for 12 h before spray inoculation. Disease symptoms were photographed at 7 d post inoculation. (B) Bacterial growth of *P. syringae* pv. *tomato* DC3000 ΔhopQ1-1 in *N. benthamiana fls2* and transgenic lines expressing AtFLS2, GbFLS2 or PtFLS2 after flg22^Pst^ pre-treatment. Bacterial populations were determined at 0, 2 and 4 d post inoculation. Data are presented as mean ± SD from three independent experiments. Different letters indicate statistically significant differences as determined by one-way ANOVA followed by Tukey ‘ s multiple-comparison test (P < 0.05). (C) Representative wilt symptoms of *N. benthamiana fls2* and transgenic lines expressing AtFLS2, GbFLS2 or PtFLS2 at 12 d after soil-drench inoculation with *Ralstonia solanacearum* strain TP2, a tobacco wilt isolate. (D) Disease index scores of bacterial wilt symptoms after soil-drench inoculation with *R. solanacearum* strain TP2. Wilt symptoms were scored on a scale from 0 to 4, where 0 = no visible symptoms and 4 = complete wilting or death. Data are presented as mean ± SD (n = 12 plants).

We next examined MAPK activation in these stable lines. Upon flg22^Pst^ treatment, AtFLS2-, GbFLS2- and PtFLS2-expressing transgenic lines all showed clear MAPK activation, whereas no phosphorylated MAPK signal was detected in the *fls2* mutant (Fig. 4C, L1; Fig. S4A, L2). Under flg22^Rso^ treatment, MAPK activation was not detected in the GbFLS2 lines, but was clearly induced in the PtFLS2 lines (Fig. 4D, L1; Fig. S4B, L2). A weak delayed MAPK signal was also detectable in the AtFLS2 line after flg22^Rso^ treatment, consistent with recent evidence that heterologous expression of AtFLS2 in *N. benthamiana* can confer responsiveness to flg22^Rso^ (Li et al., 2025). Overall, the stable transgenic lines largely recapitulated the ligand-recognition patterns observed in the transient expression assays.

We then assessed a later PTI output using a seedling growth inhibition assay. After 1 week of growth on solid medium, seedlings were transferred to liquid medium containing 5 μ M flg22^Pst^ or flg22^Rso^ and cultivated for an additional week. Under flg22^Pst^ treatment, clear growth inhibition was observed in the AtFLS2-, GbFLS2- and PtFLS2-expressing lines, whereas the *fls2* mutant remained largely insensitive. By contrast, under flg22^Rso^ treatment, obvious growth inhibition was observed only in the PtFLS2 transgenic lines, while the AtFLS2-, GbFLS2-expressing lines and the *fls2* mutant showed little or no visible response (Fig. 4E, L1; Fig. S4C, L2). Consistent with these phenotypic observations, fresh-weight measurements confirmed that flg22^Pst^ caused significant growth inhibition in the AtFLS2, GbFLS2 and PtFLS2 lines, whereas flg22^Rso^ significantly reduced fresh weight only in the PtFLS2 lines (Fig. 4F, L1; Fig. S4D, L2).

To determine whether these stable transgenic PTI outputs depend on BAK1, we silenced *NbSERK3B* in representative L1 lines expressing AtFLS2, GbFLS2 or PtFLS2 using VIGS. Silencing efficiency was confirmed by qRT-PCR (Fig. S5). Compared with the *TRV:EV* control, *NbSERK3B* silencing significantly reduced flg22^Pst^-induced ROS production in all three transgenic lines (Fig. 4G). In the PtFLS2 line, *NbSERK3B* silencing also markedly reduced the ROS response to flg22^Rso^ (Fig. 4H). Consistent with the ROS data, MAPK activation at 15 min after flg22^Pst^ treatment was attenuated in the AtFLS2, GbFLS2 and PtFLS2 lines following *NbSERK3B* silencing (Fig. 4I). Likewise, *NbSERK3B* silencing reduced flg22^Rso^-induced MAPK activation in the PtFLS2 line (Fig. 4J). Together, these results indicate that both gymnosperm FLS2 receptors remain functional in stable transgenic *N. benthamiana* and that their PTI outputs depend on BAK1 co-receptor function.

### Stable expression of gymnosperm FLS2 receptors enhances antibacterial immunity in transgenic *N. benthamiana*

We next asked whether these stable PTI outputs were associated with enhanced antibacterial immunity using representative L1 lines from each construct. To address this, we first challenged the *N. benthamiana fls2* mutant and transgenic lines expressing AtFLS2, GbFLS2 or PtFLS2 with *Pseudomonas syringae* pv. *tomato* DC3000 Δ hopQ1-1, a strain capable of causing disease in *N. benthamiana*. Under spray inoculation without peptide pre-treatment, all genotypes developed severe disease symptoms with no obvious differences among them (Fig. S6). We therefore pre-treated plants with 1 μM flg22^Pst^ for 12 h before spray inoculation. Under these conditions, disease symptoms at 7 d post inoculation were clearly reduced in the AtFLS2-, GbFLS2- and PtFLS2-expressing lines compared with the *fls2* mutant. Consistent with the disease phenotypes, bacterial growth assays showed that, following flg22^Pst^ pre-treatment, DC3000 Δ hopQ1-1 populations were significantly lower in the AtFLS2, GbFLS2 and PtFLS2 transgenic lines than in the *fls2* mutant at both 2 and 4 d post inoculation.

We next assessed resistance of the stable transgenic lines to *Ralstonia solanacearum* strain TP2, a tobacco wilt isolate, by soil-drench inoculation. At 12 d post inoculation, wilting was reduced in the AtFLS2- and PtFLS2-expressing plants relative to the *fls2* mutant and the GbFLS2 line, whereas GbFLS2 conferred only limited protection relative to the mutant background. Disease index scoring showed consistently lower wilt severity in the AtFLS2 and PtFLS2 lines than in the *fls2* mutant, whereas the GbFLS2 line showed only limited improvement. By 20 d post inoculation, all plants of the *fls2* mutant and the GbFLS2-expressing line had died, whereas surviving plants remained in the AtFLS2- and PtFLS2-expressing lines (2/12 and 4/12, respectively; Fig. S7). Thus, PtFLS2 was associated with more pronounced protection against *R. solanacearum*, whereas AtFLS2 also provided partial protection in this heterologous background.

## Discussion

### Functional conservation and diversification of gymnosperm FLS2 receptors

Since AtFLS2 was identified as the first plant pattern-recognition receptor, FLS2 has become one of the best-characterized immune receptors in plants (Gómez-Gómez & Boller, 2000; Chinchilla et al., 2006). Its broad distribution across plant lineages, together with its well-defined ligand and tractable immune outputs, has also made it a major system for studying PRR signalling mechanisms and natural variation in ligand recognition. Functional studies in angiosperms have shown that FLS2 receptors can differ substantially in ligand-recognition range. For example, tomato FLS2 recognizes the shorter *Escherichia coli*-derived flg15 peptide, whereas Arabidopsis FLS2 does not (Robatzek et al., 2007; Clarke et al., 2013). In grapevine, VvFLS2 differentially recognizes flagellin epitopes from pathogenic and beneficial bacteria, consistent with avoidance of inappropriate activation by an endophytic bacterium (Trdá et al., 2014). More recently, the wild grape receptor Vitis riparia FLS2XL was shown to perceive the flagellin of *A. tumefaciens*, and soybean GmFLS2 variants were found to recognize flg22Rso from *R. solanacearum* (Fürst et al., 2020; Wei et al., 2020). In parallel, the generation of a *N. benthamiana fls2* mutant enabled transient heterologous assays that uncovered multiple FLS2 receptors capable of perceiving Agrobacterium-derived flg22 variants (Wu et al., 2022). Together, these studies show that FLS2 is a versatile receptor system and that angiosperms contain many naturally diversified flagellin receptors. However, despite this progress, functional characterization of FLS2 has remained almost entirely confined to angiosperms, and direct experimental evidence that gymnosperms possess functional FLS2 receptors has been lacking.

Our results provide direct evidence that two gymnosperm FLS2 orthologues, GbFLS2 from G. biloba and PtFLS2 from *P. tabuliformis*, can function as bona fide flagellin receptors in a heterologous angiosperm background. Although both receptors are highly divergent from angiosperm FLS2 proteins at the sequence level, they are robustly placed within the FLS2 clade. In the *N. benthamiana fls2* mutant, both receptors restored flg22^Pst^-triggered ROS production and MAPK activation in transient assays, and both remained functional in stable transgenic plants, where they also mediated flg22^Pst^-dependent seedling growth inhibition. Importantly, our in planta cross-linking assays showed that both GbFLS2 and PtFLS2 specifically associate with flg22^Pst^, providing direct biochemical support for receptor – ligand association in planta. In the same heterologous system, early immune outputs mediated by GbFLS2 and PtFLS2 depended on NbSERK3B function, consistent with preservation of key co-receptor requirements (Heese et al., 2007; Chinchilla et al., 2007). Together, these findings suggest that the core ligand-recognition and signalling features of the FLS2 module were already present before the split between extant gymnosperms and angiosperms, although direct validation in native gymnosperm tissues will be needed to define the full extent of conservation in the endogenous context.

At the same time, conservation does not preclude diversification. As in angiosperms, our data indicate that gymnosperm FLS2 receptors can differ in ligand-recognition spectrum. GbFLS2 retained a recognition profile largely consistent with a typical FLS2 receptor, whereas PtFLS2 additionally perceived flg22^Rso^, a flagellin variant that often fails to elicit strong responses through angiosperm FLS2 receptors (Wei et al., 2020). This contrast is important because Ginkgo and Pinus represent deeply diverged gymnosperm lineages, with Ginkgo belonging to the ginkgophytes and Pinus to the conifers (Yang et al., 2022). The finding that two phylogenetically distant gymnosperm receptors already differ in flg22 recognition suggests that diversification of FLS2 function was not restricted to angiosperms and may also be distributed across gymnosperms. These results suggest that gymnosperms may contain additional FLS2 orthologues with distinct flagellin-recognition properties. Consistent with this idea, stable expression of PtFLS2 in transgenic *N. benthamiana* was associated with enhanced responsiveness to flg22^Rso^ and more pronounced resistance to *R. solanacearum*. Recent work has also shown that heterologous expression of AtFLS2 in *N. benthamiana* can confer responsiveness to flg22^Rso^ and enhance anti-bacterial immunity, consistent with the partial protection observed here in the AtFLS2-expressing lines (Li et al., 2025). Our results therefore show that expanded flg22 recognition in a heterologous system is not limited to angiosperm FLS2 receptors, but can also occur in gymnosperm FLS2 orthologues. It is worth noting that the GbFLS2 reported here was not recovered in our previous heterologous survey (Wu et al., 2022), because the earlier *G. biloba* genome assembly used in that study did not include this FLS2 sequence.

### Implications for structure-guided FLS2 analysis and ligand-binding validation

Recent studies have shown that FLS2 receptors can be identified and engineered by combining structural prediction with heterologous screening in the *N. benthamiana fls2* mutant, highlighting the growing potential of AI-assisted receptor design in plant immunity (Li et al., 2025; Zhang et al., 2025). In our study, structural modelling was used primarily to assess interface conservation and to guide experimental validation of receptor – ligand interactions, rather than to engineer new receptor variants. Our results also show that local interface features can be more informative than global confidence scores in some cases. In the GbFLS2- and PtFLS2 – flg22^Pst^ models, the interface was highly convergent across replicate predictions, recurrent hydrogen-bonding patterns were readily recovered, and mean pairwise PAE values remained low. Under these conditions, structure-guided mutagenesis was more likely to yield interpretable results. By contrast, the PtFLS2 – flg22^Rso^ models retained a relatively high overall ipTM, yet local hydrogen-bonding patterns were less reproducible and pairwise PAE values were substantially higher. In this case, residues shared with the flg22^Pst^ interface could still be validated functionally, whereas candidate flg22^Rso^-specific contacts could not be resolved with confidence. These observations suggest that, in this context, local interface convergence and pairwise PAE values may provide more useful guidance than a satisfactory global ipTM alone when prioritizing candidate residues for experimental testing. For flg22^Rso^, our data therefore support ligand perception by PtFLS2, but do not yet resolve the complete set of specificity-determining contacts.

The in planta cross-linking assay used here also helped bridge structural prediction and immune readout. This approach builds on an established methodological lineage, from early FLS2 studies in which chemical cross-linking and immunoprecipitation demonstrated direct flg22 – FLS2 association, to later peptide – receptor systems employing biotinylated ligands, GFP-Trap purification and competitive binding controls (Chinchilla et al., 2006; Albert et al., 2015; Fan et al., 2022; Burggraf & Albert, 2024). In our study, it helped distinguish mutations that abolished detectable ligand association from those that primarily compromised signalling output after ligand perception. Such assays may also be useful in future FLS2 engineering efforts, particularly in cases where direct biochemical affinity measurements remain difficult and structural models alone do not yet unambiguously define specificity-determining residues. Together, our results show that structural prediction, heterologous functional screening and ligand-association assays can be used together to analyse naturally diversified FLS2 receptors from phylogenetically distant lineages.

## Acknowledgements

This work was supported by the National Natural Science Foundation of China (grant no. 32572059).

## Author contributions

HX performed most of the experiments, participated in experimental design, and wrote the first draft of the manuscript. XH, LW and LZ participated in the experiments. QC conceived the study, developed the overall framework, and revised the manuscript.

## Data availability

The data supporting the findings of this study are available within the article and its Supporting Information. Sequence data for GbFLS2 and PtFLS2 have been deposited in GenBank under accession numbers PZ262359 and PZ262360.

