## Supporting Information (Methods, Tables and Figures) for "Gymnosperm FLS2 orthologues mediate conserved flg22 perception and divergent responses to flg22 variants"

### The following Supporting Information is available for this article:

**Methods S1 Phylogenetic analysis.**
**Methods S2 Stable transformation of *N. benthamiana fls2*.**
**Methods S3 ROS burst and MAPK activation assays.**
**Methods S4 Virus-induced gene silencing (VIGS) assays.**
**Methods S5 Site-directed mutagenesis.**
**Table S1 Primer sequences used in this study.**
**Table S2 Recurrent hydrogen-bonding contacts predicted between flg22$^{Pst}$ and GbFLS2 in 10 independent AF3 models.**
**Table S3 Recurrent hydrogen-bonding contacts predicted between flg22$^{Pst}$ and PtFLS2 in 10 independent AF3 models.**
**Table S4 Recurrent hydrogen-bonding contacts predicted between flg22$^{Rso}$ and PtFLS2 in 16 independent AF3 models.**
**Fig. S1 Domain organization of GbFLS2 and PtFLS2.**
**Fig. S2 Expanded phylogenetic analysis of GbFLS2 and PtFLS2 with angiosperm FLS2 receptors.**
**Fig. S3 ROS kinetics of PtFLS2 in response to different flg22 variants in transient assays.**
**Fig. S4 GbFLS2 and PtFLS2 mediate conserved PTI outputs in independent L2 transgenic *N. benthamiana* lines.**
**Fig. S5 qRT-PCR analysis of *NbSERK3B* silencing efficiency in VIGS-treated transgenic lines.**
**Fig. S6 Disease symptoms after spray inoculation with *Pseudomonas syringae* pv. *tomato* DC3000 ΔhopQ1-1 without flg22$^{Pst}$ pre-treatment.**
**Fig. S7 Survival of transgenic lines after inoculation with *Ralstonia solanacearum* strain TP2.**

**Methods S1 Phylogenetic analysis.**

Phylogenetic analysis of FLS2 homologues was performed using the maximum-likelihood method in MEGA 7.0 (Kumar et al., 2016) with the Jones–Taylor–Thornton (JTT) substitution model, using all aligned amino acid positions. Bootstrap support values were calculated from 1000 replicates and are indicated at the corresponding nodes. The dataset included GbFLS2 and PtFLS2 together with representative functionally validated angiosperm FLS2 receptors from monocot and dicot species. For the tree shown in Fig. 1A, four known PRRs from the RLK XII family (AtEFR, AtXPS1, SlFLS3 and StPERU) were included for comparison. For the expanded phylogenetic analysis shown in Fig. S2, additional angiosperm FLS2 sequences were selected from a previously compiled FLS2 sequence dataset (Cheng et al., 2020). Protein sequences were aligned before tree construction.

**Methods S2 Stable transformation of *N. benthamiana fls2*.**

Leaf discs excised from 4-week-old in vitro-grown *N. benthamiana fls2* seedlings were inoculated with an Agrobacterium tumefaciens suspension (OD600 = 0.5) prepared in one-quarter-strength liquid Murashige and Skoog (MS) medium for 10 min. After inoculation, the leaf discs were co-cultivated for 2 d on regeneration medium consisting of MS medium supplemented with 0.1 mg/L naphthaleneacetic acid and 1 mg/L 6-benzyladenine. The explants were then transferred to the same regeneration medium supplemented with 50 mg/L kanamycin and 200 mg/L timentin for shoot selection and bacterial suppression. Regenerated shoots were excised and transferred to rooting medium until well-developed roots had formed. T0 plants were selected on medium containing 50 mg/L kanamycin. T1 and T2 generation plants were obtained by selfing, and homozygous lines were identified on the basis of kanamycin-resistance segregation in the progeny.

**Methods S3 ROS burst and MAPK activation assays.**

Reactive oxygen species (ROS) production was measured using a luminol-based assay as described by Sang & Macho (2017). Leaf discs from the indicated *N. benthamiana* plants were floated overnight in sterile water in 96-well plates. The water was then replaced with assay solution containing 100 μM luminol, 20 μg/mL horseradish peroxidase and the indicated flg22 peptides at 200 nM. Luminescence was recorded immediately after addition of the assay solution using a GloMax™ 96 Microplate Luminometer (Promega).

For MAPK activation assays, leaf discs from the indicated *N. benthamiana* plants were floated overnight in sterile water and then treated with the indicated peptides at the indicated concentrations. Samples were collected at 0, 5 and 15 min after treatment, immediately frozen in liquid nitrogen, and used for immunoblot analysis. Phosphorylated MAPKs were detected using an anti-phosphop44/42 MAPK (Erk1/2) antibody (Thr202/Tyr204; Cell Signaling Technology) and visualized with BCIP/NBT substrate.

**Methods S4 Virus-induced gene silencing (VIGS) assays.**

Virus-induced gene silencing (VIGS) assays were performed in *N. benthamiana* using the tobacco rattle virus (TRV) vector system. *Agrobacterium tumefaciens* GV3101 cultures carrying *pTRV1* and *pTRV2:NbSERK3B* were mixed in a 1:1 ratio and infiltrated into young leaves of 3-week-old *N. benthamiana* plants. As a negative control, Agrobacterium cultures carrying *pTRV1* and the empty *pTRV2* vector were mixed in the same ratio and infiltrated in parallel. At 2 weeks after TRV infiltration, leaves at comparable developmental stages were collected for ROS burst and MAPK activation assays.

**Methods S5 Site-directed mutagenesis.**

Alanine-substitution mutants of GbFLS2 and PtFLS2 were generated by site-directed mutagenesis using the Mut Express II Fast Mutagenesis Kit (Vazyme) and verified by Sanger sequencing.

**Table S1 Primer sequences used in this study.**

| Gene or transcript name | Forward and reverse primers (5’ to 3’) | Locus or accession no./Note |
|---|---|---|
| Primers for *GbFLS2*，*PtFLS2*，*AtFLS2*，*AtEFR* cloning | | |
| *GbFLS2* | ggggactctagaataggtaccATGGCCAACCCCACCCCC<br>gcccttgctcaccatctcgagGTACTGTTCCGTCAAAGCCAAT | PZ262359 / For Gibson assembly into pH35GG-Km |
| *PtFLS2* | ggggactctagaataggtaccATGGGATCGTCGGCCTCC<br>gcccttgctcaccatctcgagATAAATATCTTCATCATGAATGCGATT | PZ262360 / For Gibson assembly into pH35GG-Km |
| *AtFLS2* | ggggactctagaataggtaccATGAAGTTACTCTCAAAGACCTTTTTGA<br>gcccttgctcaccatctcgagAACTTCTCGATCCTCGTTACGATC | AT5G46330 / For Gibson assembly into pH35GG-Km |
| *AtEFR* | ggggactctagaataggtaccATGAAGCTGTCCTTTTCACTTGTTT<br>gcccttgctcaccatctcgagCATAGTATGCATGTCCGTATTTAACATC | AT5G20480 / For Gibson assembly into pH35GG-Km |
| Primers for silencing *NbSERK3B* | | |
| Silencing fragment | cccttcacctctagacccgggTGATTGGGTGAAGGGACTCCT<br>cttgtcgacactagtaagcttGGTCGGATATTTGAAGTGGAGTCG | HQ332145 / For Gibson assembly into pTRV2 |
| Primers for qRT-PCR | | |
| *NbSERK3B* | AACCACCATTCTTGCACTTCAA<br>GATGAGGGTGTAGGAGGAAGAG | For qRT-PCR analysis of *NbSERK3B* expression |
| *NbEF1α* | AGCTTTACCTCCCAAGTCATC<br>AGAACGCCTGTCAATCTTGG | Internal control of *N. benthamiana* for qRT-PCR |

**Table S2 Recurrent hydrogen-bonding contacts predicted between flg22$^{Pst}$ and GbFLS2 in 10 independent AF3 models.**

| flg22$^{Pst}$ residue | N10 | A12 | K13 | D14 | D15 | A17 | G18 | Q20 | A22 |
|---|---|---|---|---|---|---|---|---|---|
| GbFLS2 residue | D227 / N251 | R321 | Y277 / Y301 | R321 / N345 | S325 | R417 | Y373 | S419 / T421 / S443 | D467 |
| Recurrence | 8/10, 10/10 | 7/10 | 10/10, 10/10 | 7/10, 10/10 | 10/10 | 9/10 | 9/10 | 9/10, 9/10, 9/10 | 9/10 |
| Mean atom distance (Å) | 2.90 / 3.25 | 3.16 | 3.24 / 2.82 | 3.01 / 3.07 | 2.80 | 3.01 | 3.16 | 3.01 / 2.99 / 2.80 | 2.86 |
| Mean PAE (Å) | 3.44 / 3.43 | 3.74 | 2.62 / 2.50 | 2.88 / 3.35 | 2.83 | 3.26 | 2.88 | 3.50 / 3.31 / 3.44 | 3.97 |

**Table S3 Recurrent hydrogen-bonding contacts predicted between flg22$^{Pst}$ and PtFLS2 in 10 independent AF3 models.**

| flg22$^{Pst}$ residue | N10 | A12 | D14 | A17 | G18 | Q20 | A22 |
|---|---|---|---|---|---|---|---|
| PtFLS2 residue | D223 / S225 | R317 | Y297 / R317 | K389 / Y369 | T391 | S415 / S439 | D463 |
| Recurrence | 10/10, 10/10 | 9/10 | 9/10, 9/10 | 10/10, 10/10 | 10/10 | 10/10, 10/10 | 10/10 |
| Mean atom distance (Å) | 2.59 / 3.07 | 3.23 | 4.28 / 2.60 | 2.59 / 3.97 | 2.69 | 3.02 / 2.60 | 2.69 |
| Mean PAE (Å) | 4.45 / 4.64 | 5.20 | 5.14 / 5.52 | 4.06 / 4.12 | 2.86 | 2.75 / 2.84 | 4.14 |

**Table S4 Recurrent hydrogen-bonding contacts predicted between flg22$^{Rso}$ and PtFLS2 in 16 independent AF3 models.**

| flg22$^{Rso}$ residue | R2 | R8 | S11 | A12 | D15 | S16 | A20 | S22 |
|---|---|---|---|---|---|---|---|---|
| PtFLS2 residue | D223 / D247 | D247 | R317 | R317 | K389 | H345 / Y369 | S415 / S439 | D463 / S465 |
| Recurrence | 13/16, 7/16 | 14/16 | 12/16 | 14/16 | 15/16 | 11/16, 11/16 | 15/16 | 15/16 |
| Mean atom distance (Å) | 3.33 / 2.85 | 2.72 | 3.20 | 2.39 | 2.99 | 3.26 / 3.09 | 3.14 / 2.86 | 3.05 / 2.94 |
| Mean PAE (Å) | 16.95 / 17.66 | 16.76 | 16.69 | 14.87 | 15.02 | 14.40 / 14.01 | 11.27 / 11.98 | 14.19 / 15.57 |

**(a)** **GbFLS2**

| | |
|---|---|
| | **1** |
| **SP** | MANPTPLLSSSACFLLLFTITFLHG |
| | **34** |
| **LRR-NT** | ATDVEALISFKKSISRNPLKALVD<br>WTNITHHCNWSGIACD |
| **LRR domain** | **102** |
| LRR1 | LDSLDLTSNSFHGSIPPQLALCLN |
| LRR2 | LTNLALYDNALTGTIPDSLGALVQ |
| LRR3 | LQSLDLAKNFLEGSIPESICNCTS |
| LRR4 | LTAVSFSNNNLTGAIPANIGNLLK |
| LRR5 | LQLFDAYTNDLTGSIPRSFGKCHD |
| LRR6 | LEALDLSENRLDGSIPTELGNMYS |
| LRR7 | LQYLNLFDNALSGSIPSSLSSCKN |
| LRR8 | LVQIALYKNNISGTIPGDFGKLAK |
| LRR9 | LEVLLLYQNSLYGTIPAALSLCKS |
| LRR10 | LTRLVLSNNQLTGNIPYELGSLAS |
| LRR11 | LENLSLFLNRLTGVIPASLGNCLN |
| LRR12 | LTTLALYSNNLTGPIPQSLVSLEK |
| LRR13 | LQHLLIANNELSGSIPTSLSNCSS |
| LRR14 | LVRISFTYNQLTGQIPSDLDKLSM |
| LRR15 | IRYISFGYNQLSGRIPSATYNCSL |
| LRR16 | LLILDFAGNRLTGPLQGIGKLRN |
| LRR17 | LQRLLLFLNSFSGAIPMEIGNLTN |
| LRR18 | LISLELGGNNFSGSIPASLGLLTQ |
| LRR19 | LQGLYLQKNFLDEGIPEELSGCVQ |
| LRR20 | LSELELQLNRLTGAIPEALSKLHW |
| LRR21 | LTSLYLYGNMLNGSIPSSLGKCTR |
| LRR22 | LQDLDISHNQLIGSIPTSVASLKEF |
| LRR23 | HINFNMSNNFLTGSIPTELGGMQM |
| LRR24 | VQVIDLSNNNLSGTIPSSLGECSS |
| LRR25 | LQGLDLSKNGLQGNIPVSLSRLLSL |
| LRR26 | EFFLNLSHNQLGGEIPEELTSLKL |
| LRR27 | LMSLDFSSNRLTGSIPSKLGNLTM |
| LRR28 | LKFLNLSDNQLEGPIPQVGILRNF |
| | **814** |
| **TM** | IVISVLAGITILTALISTAFIL |
| | **837** |
| **JM** | LRFKNAKKNIKPEDSMKVPAEHLT<br>RFTQRDLENATDSF |
| | **873** |
| **non-RD kinase** | NPNNIIGSSSISTVYKGVLNNGRT<br>VAIKKMSLQTSKELDRCFNTELHT<br>LARIRHRNLVKVLGFAWDNRIKAI<br>VLDFMPNGNLDALLHDVDQGGCQL<br>GFAERLQICISVAQGLVYLHEEYG<br>IPIVH**CD**LKPANILLDEDLEAHVT<br>DFGTARMLGVHLEDESGRSSSVFQ<br>GTIGYFAPEFAYMSRVTPKADVFS<br>FGIILMEILTNRRPTSNLLGGSSE<br>GGVSTLQEWIENAVRSGLSGALGV<br>VNSSLLVNSSREDEEKILGLLKIS<br>LLCTKPVPENRPSMSAVLSWLSSV<br>KQNRMNGRRSIEAEDWLLTEQY |

**(b)** **PtFLS2**

| | |
|---|---|
| | **1** |
| **SP** | MGSSASPFMVLLLLSALILEILPMYVAT |
| | **31** |
| **LRR-NT** | SEADLQGLIAFKASITSDPLNALA<br>DWTASAHHCNWSGVACD |
| **LRR domain** | **99** |
| LRR1 | LASLDLRSNFFHGAIPPQLALCSQ |
| LRR2 | LIDLELFNNSLTGTIPHTLGGLP |
| LRR3 | LQSLDLANNFLMGSIPDSICNCTS |
| LRR4 | LTAVSFSFNNLTGRIPVNIGNLVK |
| LRR5 | LQLFVAYVNELTGGIPPSFGKCTD |
| LRR6 | LEALDLSVNRLEGSIPPELGNVTS |
| LRR7 | LQYLDLFDNALTGTIPSTLSSCRN |
| LRR8 | LVQIALHKNKMTGTIPEDIGALSK |
| LRR9 | LEVLLLYTNGLSGGIPPALSHCKS |
| LRR10 | LVRLDLSDNQLTGYIAAELGSLPS |
| LRR11 | LKILHLHVNKLTGRIPPSLGRCSN |
| LRR12 | LTTLALYNNQLAGPIPQSLGLLSR |
| LRR13 | LEKLTIFSNQLSGSIPPSLFNCSS |
| LRR14 | LVNISVPNNQLTGQIPPEIGKLSM |
| LRR15 | LNFLSLGNNLLSGEVPATLYNCSL |
| LRR16 | LRKLDISRNNFSGSMKGISELKK |
| LRR17 | LELLSLHHNSYTGTVPIELGSLRN |
| LRR18 | LYSLDLGSNRFSGPFPPSLGVLTH |
| LRR19 | LQRLFLDRNYLTGRIPNEITGCRE |
| LRR20 | LAELKLDWNQFSGTIPEAISNLEW |
| LRR21 | LNLLSLSGNLLSGPIPESLRKCIR |
| LRR22 | LQNLDISYNKLTGSIPRSVASLRGL |
| LRR23 | QINFNLSNNLLSGSIPSELGGMQM |
| LRR24 | VQFVDLSNNNLSGAIPSSIAGCVG |
| LRR25 | LQGLDLSGNTLQGQIPVALSRLQNL |
| LRR26 | EQFLNLSNNHLQGEIPAELGELKL |
| LRR27 | LRSLDLSSNMLSGKIPVELGNLTS |
| LRR28 | LRTLNLSGNELEGPVPPVGILKSF |
| | **791** |
| **TM** | VVISVIVSAIISLLFCIAFILC |
| | **814** |
| **JM** | RGRRKNNINSEELSISMPGEPLIQ<br>RFTKRDLEMATES |
| | **852** |
| **non-RD kinase** | FSAGNIIGSSTISTVYRGVLRNGK<br>VVAVKILKMENSKELDQCFKRELY<br>TMARIRHRNLVKVVGYAWDNRLKA<br>LVMELMPNGNLDAVLHSVDAGEGH<br>SAQLDLSKRLSICISIAQALVYLH<br>ENYDFPIVH**CD**LKPANILLDEDFE<br>AHVSDFGTSRMLGIHLQQEASQGT<br>LSAFQLGTIGYVAPGKINLSFTLN<br>LLWIFAYMARVTPSADVFSFGIIL<br>MELLTRKRPTSFPSSSSASDQKGN<br>SLPEWIENVFKDCKSLISVIDPLL<br>LQNVSEEQEEKMTSLLKISLLCTK<br>STPENRPSMNQVLPWLLKIKENRIHDEDIY |

**Fig. S1 Domain organization of GbFLS2 and PtFLS2.**

(a) GbFLS2 and (b) PtFLS2 contain a predicted N-terminal signal peptide (SP), an LRR N-terminal domain (LRR-NT), 28 extracellular leucine-rich repeats (LRRs), a transmembrane domain (TM), a juxtamembrane region (JM) and a non-RD kinase domain. Amino acid positions marking the boundaries of major domains are indicated.

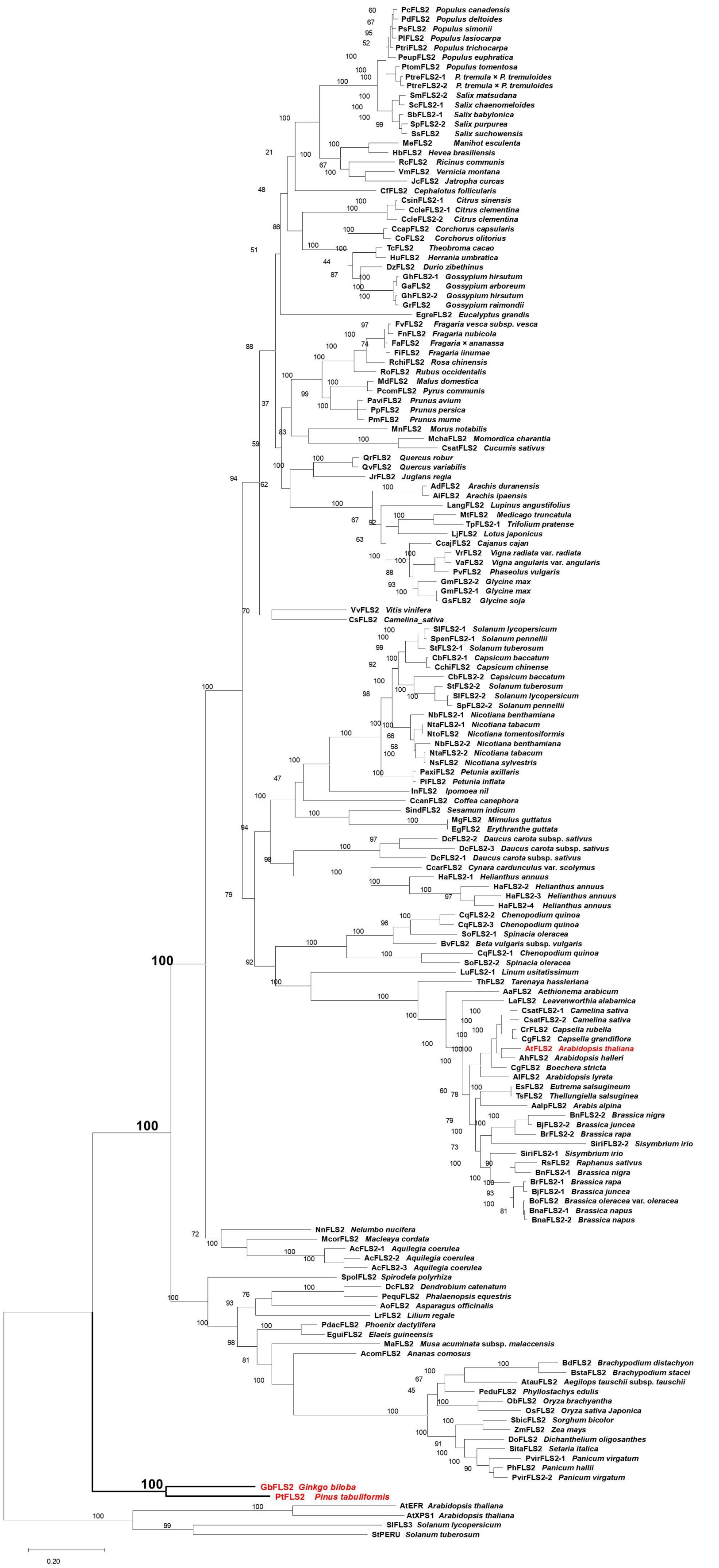
PcFLS2 Populus canadensis
PdFLS2 Populus deltoides
PsFLS2 Populus simonii
PlFLS2 Populus lasiocarpa
PtriFLS2 Populus trichocarpa
PeupFLS2 Populus euphratica
PtomFLS2 Populus tomentosa
PtreFLS2-1 P. tremula × P. tremuloides
PtreFLS2-2 P. tremula × P. tremuloides
SmFLS2-2 Salix matsudana
ScFLS2-1 Salix chaenomeloides
SbFLS2-1 Salix babylonica
SpFLS2-2 Salix purpurea
SsFLS2 Salix suchowensis
MeFLS2 Manihot esculenta
HbFLS2 Hevea brasiliensis
RcFLS2 Ricinus communis
VmFLS2 Vernicia montana
JcFLS2 Jatropha curcas
CfFLS2 Cephalotus follicularis
CsinFLS2-1 Citrus sinensis
CcleFLS2-1 Citrus clementina
CcleFLS2-2 Citrus clementina
CcapFLS2 Corchorus capsularis
CoFLS2 Corchorus olitorius
TcFLS2 Theobroma cacao
HuFLS2 Herrania umbratica
DzFLS2 Durio zibethinus
GhFLS2-1 Gossypium hirsutum
GaFLS2 Gossypium arboreum
GhFLS2-2 Gossypium hirsutum
GrFLS2 Gossypium raimondii
EgreFLS2 Eucalyptus grandis
FvFLS2 Fragaria vesca subsp. vesca
FnFLS2 Fragaria nubicola
FaFLS2 Fragaria × ananassa
FiFLS2 Fragaria iinumae
RchiFLS2 Rosa chinensis
RoFLS2 Rubus occidentalis
MdFLS2 Malus domestica
PcomFLS2 Pyrus communis
PaviFLS2 Prunus avium
PpFLS2 Prunus persica
PmFLS2 Prunus mume
MnFLS2 Morus notabilis
MchaFLS2 Momordica charantia
CsatFLS2 Cucumis sativus
QrFLS2 Quercus robur
QvFLS2 Quercus variabilis
JrFLS2 Juglans regia
AdFLS2 Arachis duranensis
AiFLS2 Arachis ipaensis
LangFLS2 Lupinus angustifolius
MtFLS2 Medicago truncatula
TpFLS2-1 Trifolium pratense
LjFLS2 Lotus japonicus
CcajFLS2 Cajanus cajan
VrFLS2 Vigna radiata var. radiata
VaFLS2 Vigna angularis var. angularis
PvFLS2 Phaseolus vulgaris
GmFLS2-2 Glycine max
GmFLS2-1 Glycine max
GsFLS2 Glycine soja
VvFLS2 Vitis vinifera
CsFLS2 Camelina_sativa
SlFLS2-1 Solanum lycopersicum
SpenFLS2-1 Solanum pennellii
StFLS2-1 Solanum tuberosum
CbFLS2-1 Capsicum baccatum
CchiFLS2 Capsicum chinense
CbFLS2-2 Capsicum baccatum
StFLS2-2 Solanum tuberosum
SlFLS2-2 Solanum lycopersicum
SpFLS2-2 Solanum pennellii
NbFLS2-1 Nicotiana benthamiana
NtaFLS2-1 Nicotiana tabacum
NtoFLS2 Nicotiana tomentosiformis
NbFLS2-2 Nicotiana benthamiana
NtaFLS2-2 Nicotiana tabacum
NsFLS2 Nicotiana sylvestris
PaxiFLS2 Petunia axillaris
PiFLS2 Petunia inflata
InFLS2 Ipomoea nil
CcanFLS2 Coffea canephora
SindFLS2 Sesamum indicum
MgFLS2 Mimulus guttatus
EgFLS2 Erythranthe guttata
DcFLS2-2 Daucus carota subsp. sativus
DcFLS2-3 Daucus carota subsp. sativus
DcFLS2-1 Daucus carota subsp. sativus
CcarFLS2 Cynara cardunculus var. scolymus
HaFLS2-1 Helianthus annuus
HaFLS2-2 Helianthus annuus
HaFLS2-3 Helianthus annuus
HaFLS2-4 Helianthus annuus
CqFLS2-2 Chenopodium quinoa
CqFLS2-3 Chenopodium quinoa
SoFLS2-1 Spinacia oleracea
BvFLS2 Beta vulgaris subsp. vulgaris
CqFLS2-1 Chenopodium quinoa
SoFLS2-2 Spinacia oleracea
LuFLS2-1 Linum usitatissimum
ThFLS2 Tarenaya hassleriana
AaFLS2 Aethionema arabicum
LaFLS2 Leavenworthia alabamica
CsatFLS2-1 Camelina sativa
CsatFLS2-2 Camelina sativa
CrFLS2 Capsella rubella
CgFLS2 Capsella grandiflora
AtFLS2 Arabidopsis thaliana
AhFLS2 Arabidopsis halleri
CgFLS2 Boechera stricta
AlFLS2 Arabidopsis lyrata
EsFLS2 Eutrema salsugineum
TsFLS2 Thellungiella salsuginea
AalpFLS2 Arabis alpina
BnFLS2-2 Brassica nigra
BjFLS2-2 Brassica juncea
BrFLS2-2 Brassica rapa
SiriFLS2-2 Sisymbrium irio
SiriFLS2-1 Sisymbrium irio
RsFLS2 Raphanus sativus
BnFLS2-1 Brassica nigra
BrFLS2-1 Brassica rapa
BjFLS2-1 Brassica juncea
BoFLS2 Brassica oleracea var. oleracea
BnaFLS2-1 Brassica napus
BnaFLS2-2 Brassica napus
NnFLS2 Nelumbo nucifera
McorFLS2 Macleaya cordata
AcFLS2-1 Aquilegia coerulea
AcFLS2-2 Aquilegia coerulea
AcFLS2-3 Aquilegia coerulea
SpolFLS2 Spirodela polyrhiza
DcFLS2 Dendrobium catenatum
PequFLS2 Phalaenopsis equestris
AoFLS2 Asparagus officinalis
LrFLS2 Lilium regale
PdacFLS2 Phoenix dactylifera
EguiFLS2 Elaeis guineensis
MaFLS2 Musa acuminata subsp. malaccensis
AcomFLS2 Ananas comosus
BdFLS2 Brachypodium distachyon
BstaFLS2 Brachypodium stacei
AtauFLS2 Aegilops tauschii subsp. tauschii
PeduFLS2 Phyllostachys edulis
ObFLS2 Oryza brachyantha
OsFLS2 Oryza sativa Japonica
SbicFLS2 Sorghum bicolor
ZmFLS2 Zea mays
DoFLS2 Dichanthelium oligosanthes
SitaFLS2 Setaria italica
PvirFLS2-1 Panicum virgatum
PhFLS2 Panicum hallii
PvirFLS2-2 Panicum virgatum
GbFLS2 Ginkgo biloba
PtFLS2 Pinus tabuliformis
AtEFR Arabidopsis thaliana
AtXPS1 Arabidopsis thaliana
SlFLS3 Solanum lycopersicum
StPERU Solanum tuberosum
0.20

**Fig. S2 Expanded phylogenetic analysis of GbFLS2 and PtFLS2 with angiosperm FLS2 receptors.**

Maximum-likelihood phylogenetic tree including GbFLS2, PtFLS2 and 112 representative angiosperm FLS2 proteins. Bootstrap support values were calculated from 1000 replicates and are shown at the corresponding nodes. GbFLS2 and PtFLS2 are highlighted in red. Scale bar, substitutions per site.

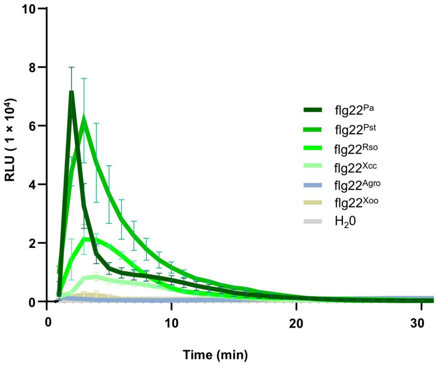


**Fig. S3 ROS kinetics of PtFLS2 in response to different flg22 variants in transient assays.**

Reactive oxygen species (ROS) production was measured in *N. benthamiana fls2* leaf discs transiently expressing PtFLS2 after treatment with the indicated flg22 peptides at 200 nM. Data are presented as mean ± SD (n = 6 leaf discs). Data are shown from one representative experiment repeated three times with similar results.

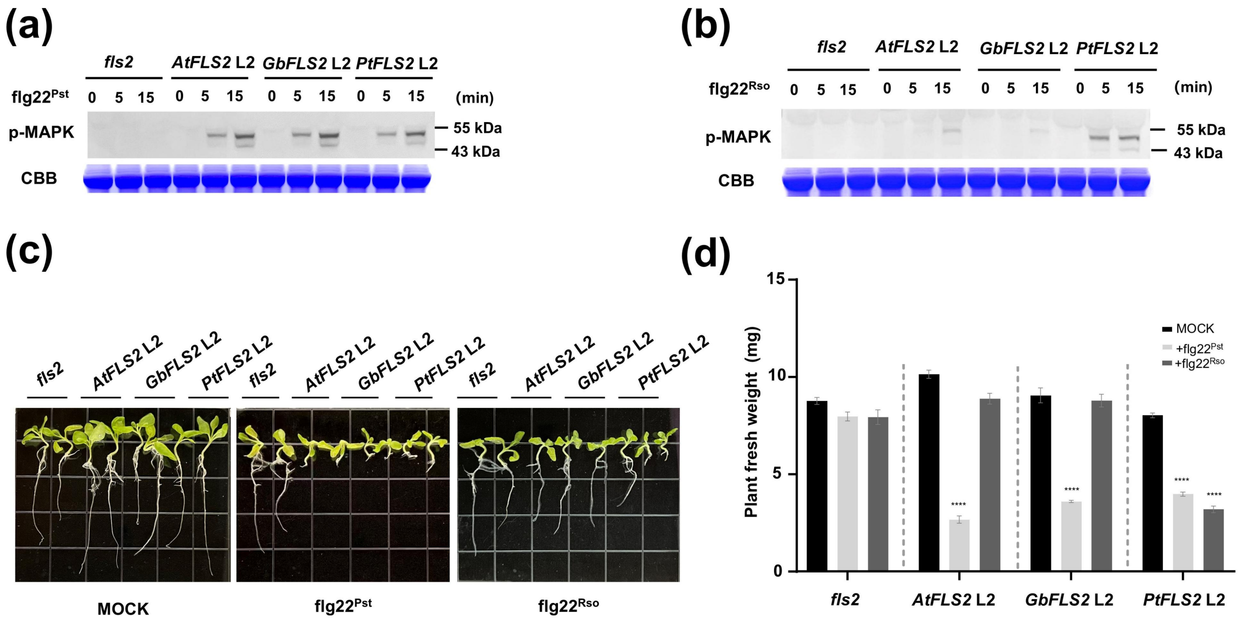


**Fig. S4 GbFLS2 and PtFLS2 mediate conserved PTI outputs in independent L2 transgenic *N. benthamiana* lines.**

(A, B) flg22-induced MAPK activation in *N. benthamiana fls2* and independent L2 transgenic lines expressing AtFLS2, GbFLS2 or PtFLS2. Leaf discs were treated with 1 μM flg22$^{Pst}$ (A) or 1 μM flg22$^{Rso}$ (B) for the indicated times, and phosphorylated MAPKs were detected by immunoblot using an anti-phospho-p44/42 MAPK antibody. Coomassie brilliant blue (CBB) staining is shown as a loading control.

(C) Seedling growth inhibition assay in *N. benthamiana fls2* and independent L2 transgenic lines. Seedlings grown on solid medium for 1 week were transferred to liquid medium containing 5 μM flg22$^{Pst}$, 5 μM flg22$^{Rso}$ or mock medium without peptide, and cultivated for an additional week before imaging.

(D) Fresh-weight measurements corresponding to the seedling growth inhibition assay shown in (C). Data are presented as mean ± SD (n = 12 seedlings). Asterisks indicate significant differences relative to the corresponding mock treatment as determined by Student's t-test (****, $P < 0.0001$).

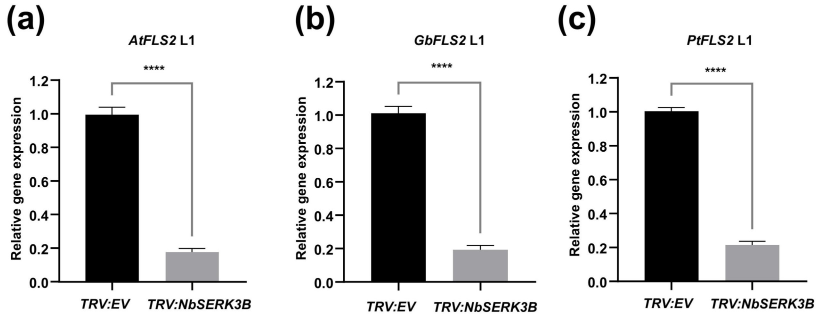


**Fig. S5 qRT-PCR analysis of *NbSERK3B* silencing efficiency in VIGS-treated transgenic lines.** Relative transcript levels of *NbSERK3B* were determined by qRT-PCR in representative L1 transgenic *N. benthamiana* lines expressing (a) AtFLS2, (b) GbFLS2 or (c) PtFLS2 after *TRV:EV* or *TRV:NbSERK3B* treatment. Expression levels were normalized to *NbEF1α*. Data are presented as mean ± SD (n = 3). Asterisks indicate significant differences as determined by Student's *t*-test (****, $P < 0.0001$).

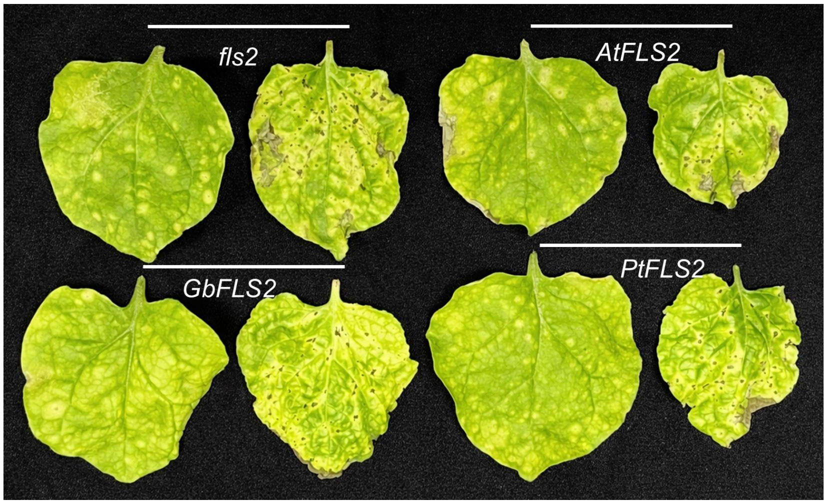


**Fig. S6 Disease symptoms after spray inoculation with *Pseudomonas syringae* pv. *tomato* DC3000 ΔhopQ1-1 without flg22$^{Pst}$ pre-treatment.**

Representative disease symptoms in *N. benthamiana fls2* and transgenic lines expressing AtFLS2, GbFLS2 or PtFLS2 at 7 d after spray inoculation without flg22$^{Pst}$ pre-treatment. Under these conditions, all genotypes developed severe disease symptoms with no obvious differences.

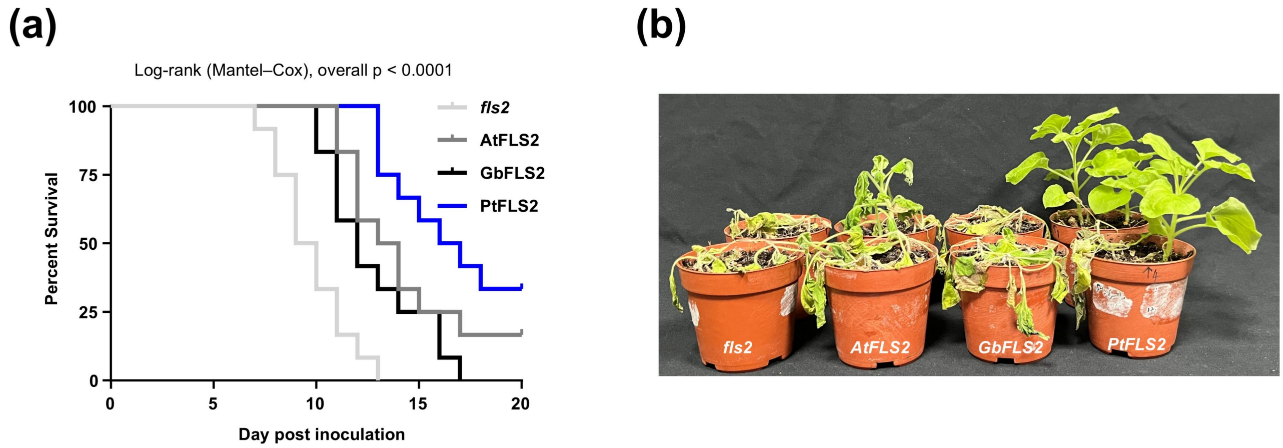


**Fig. S7 Survival of transgenic lines after inoculation with *Ralstonia solanacearum* strain TP2.**
(A) Kaplan–Meier survival curves of *N. benthamiana fls2* and representative transgenic lines expressing AtFLS2, GbFLS2 or PtFLS2 after soil-drench inoculation with *R. solanacearum* strain TP2. Overall differences among groups were analysed using the log-rank (Mantel–Cox) test.
(B) Representative disease symptoms of the indicated lines at 20 d post inoculation with *R. solanacearum* strain TP2.
